# Bottleneck Size-Dependent Changes in the Genetic Diversity and Specific Growth Rate of a Rotavirus A Strain

**DOI:** 10.1101/702233

**Authors:** Syun-suke Kadoya, Syun-ichi Urayama, Takuro Nunoura, Miho Hirai, Yoshihiro Takaki, Masaaki Kitajima, Toyoko Nakagomi, Osamu Nakagomi, Satoshi Okabe, Osamu Nishimura, Daisuke Sano

## Abstract

RNA viruses form a dynamic distribution of mutant swarm (termed “quasispecies”) due to the accumulation of mutations in the viral genome. The genetic diversity of a viral population is affected by several factors, including a bottleneck effect. Human-to-human transmission exemplifies a bottleneck effect in that only part of a viral population can reach the next susceptible hosts. In the present study, the rhesus rotavirus (RRV) strain of Rotavirus A was serially passaged five times at a multiplicity of infection (MOI) of 0.1 or 0.001 in duplicate (the 1st and 2nd lineages), and three phenotypes (infectious titer, cell binding ability and specific growth rate) were used to evaluate the impact of a bottleneck effect on the RRV population. The specific growth rate values of lineages passaged under the stronger bottleneck (MOI of 0.001) were higher after five passages. The nucleotide diversity also increased, which indicated that the mutant swarms of the lineages under the stronger bottleneck effect were expanded through the serial passages. The random distribution of synonymous and non-synonymous substitutions on rotaviral genome segments indicated that almost all mutations were selectively neutral. Simple simulations revealed that the presence of minor mutants could influence the specific growth rate of a population in a mutant frequency-dependent manner. These results indicate that a stronger bottleneck effect can create more sequence spaces for minor mutants originally existing in a hidden layer of mutant swarm.

**IMPORTANCE:** In this study, we investigated a bottleneck effect on an RRV population, which may drastically impact a viral population structure. RRV populations were serially passaged under two levels of a bottleneck effect, which exemplified a human-to-human transmission. As a result, the genetic diversity and specific growth rate of RRV populations increased under the stronger bottleneck effect, which implied that a bottleneck could create a new sequence space in a population for minor mutants originally existing in a hidden layer of a mutant swarm of the double-stranded RNA virus. The results of this study suggest that the genetic drift caused by a bottleneck in a human-to-human transmission explains the random appearance of new genetic lineages causing viral outbreaks, which can be expected by the molecular epidemiology using next generation sequencing in which the viral genetic diversity within a viral population is investigated.

## INTRODUCTION

RNA viruses form a dynamic distribution of a mutant swarm, termed quasispecies (1, 2), because a proofreading function is absent in RNA-dependent RNA polymerase (RdRp). Such dynamic genetic diversity of RNA viruses is advantageous for better adaptation to a given environment. For example, the replicative ability of West Nile virus has increased with an expansion of the size of a mutant swarm (3), and minor mutants of rabies virus have affected the adaptation to a new host even when a master sequence was not replaced (4). These examples demonstrate that the increase of genetic diversity can provide phenotypic benefits to a virus population.

Infinite incrementation of genetic diversity within an RNA virus population is restricted by external factors in an environment, such as natural selection (advantageous genes are fixed in the population or deleterious mutations are removed from the population) and bottleneck effect (genes accidentally fix in and disappear from a population when the size of that population decreases temporarily) (5). Such a bottleneck effect may impact the viral population structure, and a transmission from host-to-host (even within a host) can work on a virus population as a bottleneck effect (6, 7). For example, Vigunuzzi *et al.* reported that the diversity of a mutant swarm determined the poliovirus pathogenesis as the reduced diversity of poliovirus led to the loss of neurotropism and the attenuated pathogenicity (8). In a recent study, hepatitis C virus was serially passaged more than 50 times with a small bottleneck (about 10-fold dilution), which resulted in a decrease in drug sensitivity along with an increase in genetic diversity (9). Understanding the transition of a viral population structure during host-to-host transmissions may help us determine the implications of the occurrence of virulent viral variants in the future.

In this study, we investigated a change in population structure during serial passages (bottleneck events) using rhesus rotavirus (RRV) as a model virus. Rotavirus is a double-stranded RNA (dsRNA) virus with 11 genome segments that is transmitted via the fe-cal-oral route and causes severe diarrhea among children under five years old. Despite recommendations to vaccinate and improve hygiene (10), rotavirus outbreaks have occurred sporadically both in developed and developing countries (11–14). Assuming a human-to-human transmission, RRV was serially passaged under weaker or stronger bottleneck events (multiplicity of infection (MOI) of 0.1 or 0.001, respectively) without any positive selections. During serial passages, the changes of three phenotypes (infectious titer, a cell binding ability and a specific growth rate) were confirmed. We then estimated the nucleotide diversity of 11 genome segments as an indicator of genetic diversity, and a rank abundance analysis was performed to confirm whether a mutant swarm was expanded or reduced using the single nucleotide polymorphisms (SNPs) information. The synonymous (dS) and non-synonymous (dN) substitution rates were also calculated to confirm the existence of selective pressures other than negative selection for mutants lacking infective or replicative ability. Finally, the mutation rates per cycle of 11 genome segments were estimated using the BEAST2 software package (15). Based on these results, the relationship between phenotypes and genetic diversity of an RRV population is discussed.

## MATERIALS AND METHOD

### Virus and cell

MA104 cell lines (ATCC: CRL-2378.1™) were grown in Eagle’s MEM containing 10% fetal bovine serum (FBS), 2 mM L-Glutamine, 1% Penicillin Streptomycin (GIBCO by Life Technology) and 1.125 g/L sodium bicarbonate (Wako Pure Chemical Industries, Ltd.) in a T75 flask. Before the serial passages, RRV (genotype G3P[3]) was propagated three times on MA104 cell monolayers with serum-free Eagle’s MEM to adapt to our laboratory experimental condition.

### Serial passages

RRV suspension (10^6-7^ PFU/mL) was diluted with serum-free MEM to adjust the MOI of 0.1 or 0.001, and 4 μL of 1 μg/μL trypsin from porcine pancreas was added. This mixed suspension in a 1.5 mL tube was put in an incubator at 37 °C with 5 % CO_2_ for 30 min. After incubation, the medium in a T75 flask was removed and washed twice with Dulbecco’s PBS (−) (Nissui Pharmaceutical Co., Ltd.), and then 1mL of RRV suspension was inoculated onto the confluent MA104 cells. The flask was incubated at 37 °C with 5 % CO_2_ for 60 min. After incubation, 32 mL of serum-free Eagle’s MEM was added to the MA104 cells, and re-incubated at 37 °C with 5 % CO_2_ for 2 or 3 days. The freeze-melting cycle was done three times after incubation. The suspension that had been moved into a 50 mL tube from the flask was centrifuged at 12600×g for 10 min at 4 °C and filtered with a 0.2 μm filter to remove the cell fractions. The collected RRV suspension was inoculated to the new MA104 cell again. The RRV population was passaged five times (named as cycles 0, 1, 2, 3, 4 and 5), and this series of experiments was repeated twice for reproducibility (the 1st and 2nd lineages). Let us denote the RRV populations obtained from the 1st and 2nd serial passages at MOI of 0.1 by 0.1MOI-1 and 0.1MOI-2, and those at MOI of 0.001 by 0.001MOI-1 and 0.001MOI-2. The number following each population code name is the passage number, id est, 0.1MOI-1_5 is the 1st population lineage passaged five times at MOI of 0.1.

The infectious titer (PFU/mL) of each passage was measured by plaque assay. According to our previous report (16), serially diluted RRV suspensions were treated with 1 μg/μL trypsin from a porcine pancreas. Then, the RRV suspensions were inoculated onto confluent MA104 cells in the 6-well plate, and the plate was incubated at 37 °C with 5 % CO_2_ for 90 min. After incubation, the virus inoculate was removed, and the cells were washed twice with Dulbecco’s PBS (−) and then overlaid with 2.5 % agar, including 2 % FBS, 2% Penicillin Streptomycin, 4 mM L-Glutamine, 2.25 g/L NaHCO_3_ and 4 μg/mL trypsin from porcine pancreas. The 6-well plates were incubated for 2 days and then dyed with 0.015 % neutral red for 3 hours. After 1 or 2 days, the plaque numbers were counted.

### Cell binding assay

A cell-binding assay was conducted according to the steps outlined in previous reports (16, 17). MA104 cells were incubated on the 24 well-plate for 2 or 3 days, and then the cell culture medium was removed and washed twice with tris-buffered saline (TBS). The 100 μL of suspensions containing RRV (MOI of 1.0) were inoculated to each well and incubated at 4 °C for 1 hour with agitation every 15 min. After incubating and removing the virus suspension, the cells were washed three times with TBS, and then 140 μL of Dulbecco’s PBS and 560 μL of RNA extraction buffer (Buffer AVL of QIAamp Viral RNA Mini Kit: Qiagen, Germany) were added to each well of the plate. According to the manufacturer’s protocol, dsRNA was then purified and recovered. The extracted dsRNA was denatured at 95°C for 5 min, and placed on ice for 2 min. The denatured dsRNA (single-stranded RNA) was reverse transcribed to cDNA using the PrimeScript RT Reagent Kit (Perfect Real Time) (Takara Bio Inc., Kusatsu, Japan) according to the manufacturer’s protocol. The amount of cDNA was quantified by quantitative PCR (qPCR). The qPCR was performed with Premix ExTaq (Perfect Real Time) and a Taqman probe in an Applied Biosystems 7500 Real Time PCR System. The sequences of forward and reverse primers targeted to the NSP3 region were suggested by Pang *et al.* (Forward: 5’-/ACCATCTACACATGACCCTC/-3’, Reverse: 5’-/GGTCACATAACGCCCC/-3’) (18). The sequence of the Taqman probe suggested by Pang *et al.* was modified slightly (5’-/56-FAM/ATGAGCACA/ZEN/ATAGTTAAAAGCTAACACTGTCAA/-3’). The qPCR amplification was performed with 1 cycle of an initial denaturation step at 95°C for 5 min followed by 45 cycles of 94 °C for 20 sec and 60 °C for 1 min. Finally, 1 cycle of 72 °C for 5 min was conducted for extension.

### Specific growth rate

Following the recommendations from our previous report (16), MA104 cells were inoculated on the T25 flask for 2 - 3 days, and then the cell culture medium was removed and washed twice with Dulbecco’s PBS. The 400 μL of suspensions containing RRV (MOI of 0.01) were inoculated to the flasks individually. The supernatants were taken after 0, 6, 12, 18, 24 and 36 hours, and then infectious titer was measured by the plaque assay as described above. For estimating the specific growth rate, the modified Gompertz model was applied to the data set (19):

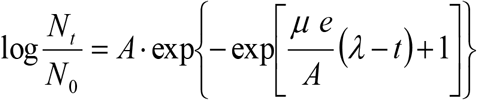

where *N*_*0*_ and *N*_*t*_ are the virus titer values (PFU/mL) at 0 and *t*-hour post-inoculation, *A* is the asymptotic value [*log*(*N*_∞_/*N*_*0*_)], *μ* is the specific growth rate [hour^−1^], *e* is the Napier’s constant and *λ* is the lag period [hour]. These parameters were determined by the least-squares method.

### Fragmented and primer ligated dsRNA sequencing (FLDS)

FLDS, a method of cDNA library construction and sequencing for dsRNA, was utilized to obtain the high sequence coverage of RRV genome sequences compared to the conventional RNA-seq (20, 21). Briefly, the total nucleic acid in each sample of serial passages was extracted by phenol-chloroform (pH 5.2), and the extracted dsRNA was purified by cellulose D (ADVANTEC, Tokyo, Japan). The dsRNA was fragmented physically by Covaris S220 (Woburn, MA, USA) and then reverse transcribed by SMARTer RACE 5’/3’ kit (Takara Bio, Inc.). The cDNA was amplified using KOD-Plus neo (Toyobo Co. Ltd., XX, Japan). Small-size cDNA and primer dimer were removed by SPRI select (Beckman Coulter, Inc.). Then the cDNA, which was fragmented and aligned with 400 base pairs (bp), was ligated with an adapter (KAPA Biosystems, Woburn, MA, USA) for Illumina MiSeq. The concentration of cDNA library was quantified by a KAPA Library Quantification Kit (KAPA Biosystems). Each 300 bp of the paired-end sequences was determined by Illumina MiSeq (Illumina Inc, San Diego, CA, USA).

MiSeq sequencing reads underwent adaptor clipping and quality trimming using Trimmomatic v0.33 (22). PhiX sequences derived from control libraries and experimentally contaminated sequences were also removed using a mapping tool, Bowtie2 ver. 2.2.9 (23). Primer sequences used for cDNA synthesis and amplification were trimmed with Cutadapt ver. 1.15 (24). Low complexity reads and PCR duplicates were detected and removed with PRINSEQ ver. 0.20.4 (25). The processed reads were used for mapping to the reference genome and generating the consensus sequence by the CLC Genomics Workbench ver.11 (CLC Bio, Aarhus, Denmark). Reference sequences of each RRV genome segment were obtained from NCBI (https://www.ncbi.nlm.nih.gov/) using the following parameters: mismatch cost of 2, insertion/deletion cost of 3, length fraction of 0.8 and similarity fraction of 0.8. Using the function “Extract consensus sequences”, the consensus sequence was extracted from the population before serial passages (cycle 0) in order to utilize it as a reference sequence. Sequence reads in each population were mapped to the cycle 0 reference sequence (“Map reads to reference”) and then realigned locally to modify the gap generated in mapping to the reference sequences (“Local realignments”).

### Estimation of nucleotide diversity, synonymous (dS) /non-synonymous (dN) substitution rate

The frequency of single nucleotide polymorphisms (SNPs) was calculated by “Low Frequency Variant Detection,” which is a function of CLC Genomics Workbench. All SNPs on each genome segment were displayed using the “circlize” of the R software package (26). In order to confirm whether the observed SNPs were the synonymous or non-synonymous amino acid substitutions, the “Amino Acid Changes” function of CLC Genomics Workbench was used. The software SNPGenie, developed by Nelson and Hughes (27) and based on the Nei-Gojobori Method (28), was used to calculate the nucleotide diversity, dN and dS.

### Rank-abundance analysis

To visualize the change in the mutant swarm of each lineage, the SNPs were arranged according to the rank based on the frequency using the R software package of “RADanalysis” (https://CRAN.R-project.org/package=RADanalysis).

### Mutation rate per cycle

By applying NEXUS files containing SNPs sequences aligned by MEGA7 (29), mutation rates of 11 genome segments were inferred using the BEAST2 software (http://www.beast2.org/) based on the Bayesian Markov chain Monte Carlo (MCMC) method (15). A GTR+γ model as the substitution site model and a Coalescent Bayesian Skyline plot for the coalescent model were applied. Under a strict molecular clock, the BEAST software was run, and then the output files were analyzed using TRACER software (30).

### Simulation of the effect of minor mutants on the specific growth rate of a population

The effect of minor mutants (or subpopulations) on a specific growth rate of a population was simulated using the modified Gompertz model (19). The growth simulations under the following four scenarios were conducted: *Simulation 1*: the frequency of dominant strain was 99 %, whereas that of minor mutants was 1 %. The specific growth rate of the dominant strain was set as 0.5 (h^−1^) (common among all four scenarios), whereas that of minor mutants was set as 2.0 (h^−1^). *Simulation 2*: the frequency of minor mutants was set as 10 %. *Simulation 3*: we focused on the VP7 genome segment of the 0.001MOI-1_5 population. Nineteen replacements per 1000 bp on VP7 genome segment can be found, and we assumed that 19 independent subpopulations possessing each nucleotide replacement were included in a population. The frequency of 19 subpopulations was 1.0 % each (in total 19%). *Simulation 4*: the frequency of minor mutant was 1 %, but the specific growth rate was 5.0 (h^−1^). The sum of the population size of a dominant strain and of minor mutants was regarded as the total population. The growth curve of the total population at each scenario was used to estimate the specific growth rate of the total population using the modified Gompertz model, under the assumption that the lag period (*λ* = 6.0 [h]) and the asymptote (*A* = 3.5 [*log N*_∞_/*N*_*0*_]) were identical among all the strains.

### Statistical analysis

A Student’s t-test was performed for infectious titer, cell binding ability and specific growth rate between the original and the cycle-5 populations. ANOVA was conducted to confirm whether a stronger bottleneck affected the nucleotide diversity. All statistical tests were performed using R software ver 3.5.0 (https://www.r-project.org/).

### Nucleotide sequence accession numbers

Rotavirus sequence data from illumina Miseq were deposited in the DDBJ databases under the accession number (DRA006847 and DRA008653).

## RESULTS

### Infectious titer, cell binding ability and specific growth rate

RRV populations were serially passaged five times at different MOI values (0.1 or 0.001). Infectious titer ranged from 10^6^ to 10^7.5^ PFU/mL, and only the 0.1MOI-1_2, 0.1MOI-1_3, and 0.1MOI-1_5 showed significant increases in infectious titer from the original population (p-value: 0.015, 0.006, 0.007, respectively) (Figure 1a). The cell binding ability was indicated by the binding efficiency to cells (the proportion of genome copies from virions bound to cells to those in the inoculum) (Figure 1b). Fewer virions (less than 2 %) were able to bind to cells, but no statistical changes were observed during the serial passages. The specific growth rate was derived from the modified Gompertz model by fitting the time-course data of infectious titer (Figure 1c). Average (± standard deviation) specific growth rates of 0.1MOI-1_5 (0.25 ± 0.004 [h^−1^]) and 0.1MOI-2_5 (0.20 ± 0.02 [h^−1^]) were not different from that of the initial populations (1st lineage: 0.22 ± 0.02, 2nd lineage: 0.25 ± 0.02 [h^−1^]). On the other hand, the specific growth rates of 0.001MOI-1_5 (1.08 ± 0.16 [h^−1^]) and 0.001MOI-2_5 (0.84 ± 0.17 [h^−1^]) were significantly higher than those of each initial population (p < 0.05).

**Figure 1.**
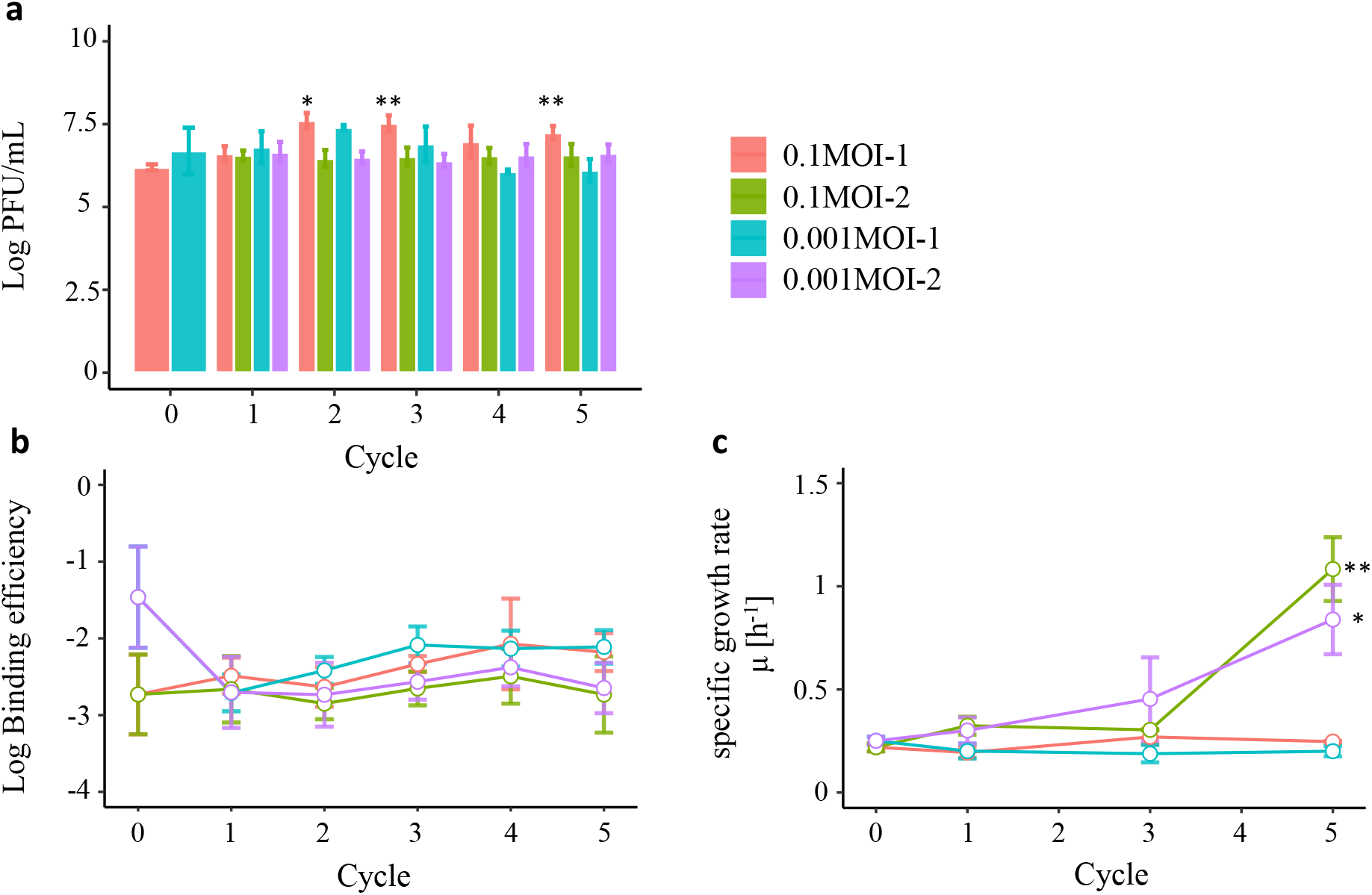
Infectious titer, cell-binding ability and specific growth rate. PFU/mL as infectious titer (a), host cell-binding efficiency (b) and specific growth rate (c) were estimated to confirm whether the serial passages altered the RRV phenotypes. All experiments were conducted three times. The error bar indicates the standard deviation. No significant differences in cell-binding ability were observed. There were significant differences in infectious titer of 0.1 MOI-1_2, 3 and 5 populations from its original population. The specific growth rates of 0.001 MOI-1_5 and 0.001 MOI-2_5 populations were also significantly larger than the original population (**: p < 0.01, *: p < 0.05, Student’s t-test).

### Nucleotide diversity of RRV populations

Using the frequency of SNPs, the nucleotide diversity of each genome segment was calculated as the indicator of genetic diversity (Figure 2). Nucleotide diversity is defined as the proportion of nucleotide difference observed when two copies of genome are sampled. Among the 0.001MOI lineages, the increments of nucleotide diversity after the cycle 4 were seen in VP1, VP2, VP3, VP6, VP7, NSP3, NSP4 and NSP5/6 genome segments. ANOVA was then conducted to investigate the effect of the MOI on nucleotide diversity. As a result, except for the VP1 genome segment (p = 0.058), significant differences in the effect on nucleotide diversity between the MOI of 0.1 and 0.001 were observed.

**Figure 2.**
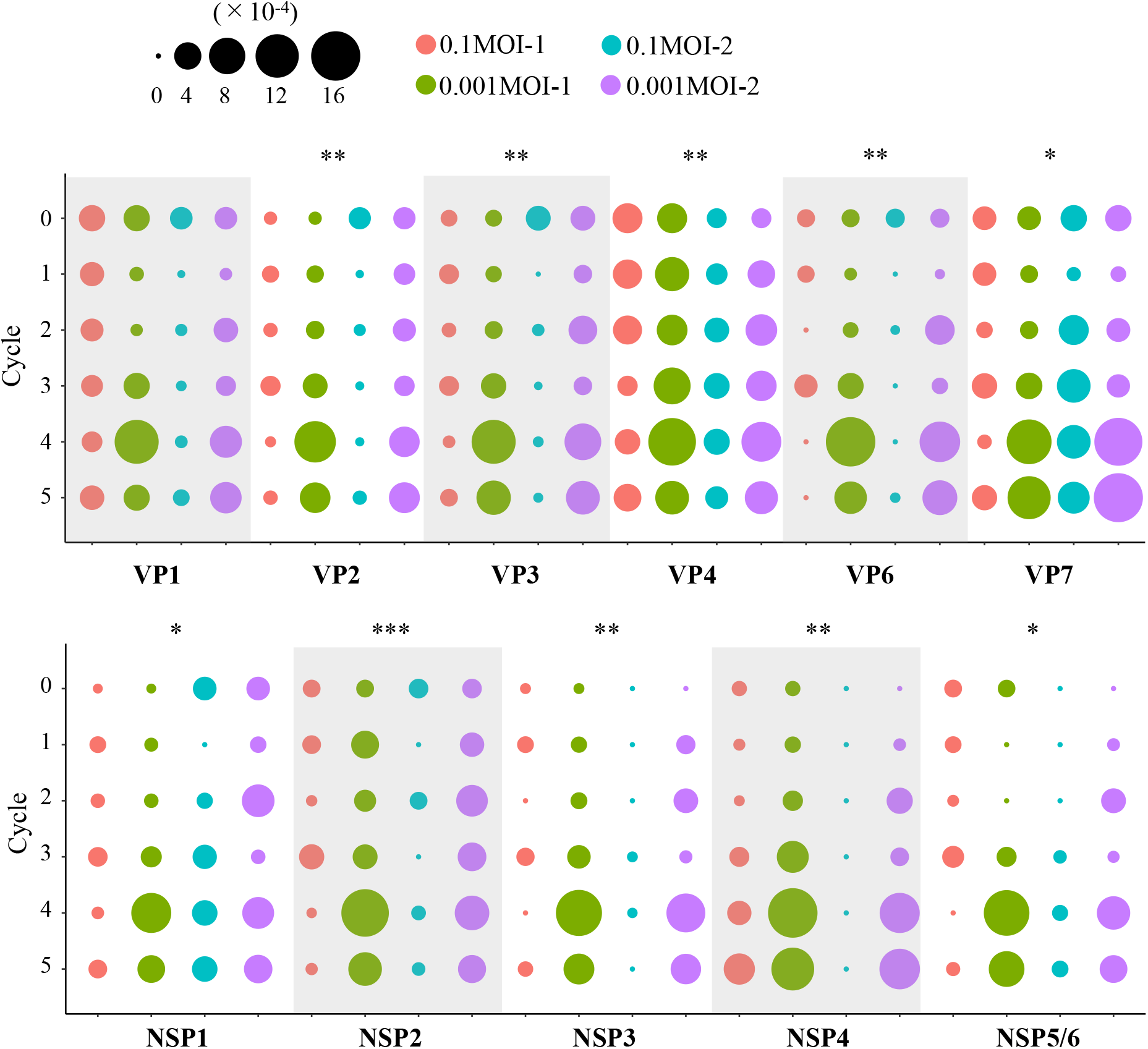
Genetic diversity of rhesus rotavirus populations. Transition of nucleotide diversity of all RRV populations is displayed as a bubble chart. The size of the plot corresponds to the value of nucleotide diversity (from 0 to 1.6×10^−3^). ANOVA was conducted to investigate the effect of the MOI on nucleotide diversity (*: p-value < 0.5, **: p-value < 0.01, ***: p-value <0.001).

### Rank abundance distribution

A rank abundance distribution of the 0.1MOI-1 lineage did not display a large change through serial passages (Figure 3). For example, the maximum rank of the 0.1MOI-2 lineage was only around 20, which implied a smaller mutant swarm whereas both 0.001MOI lineages depicted the milder sloping distributions from rank 10 or less, and the rank reached more than 100. These results indicated that mutant swarms of 0.001MOI lineages expanded during the serial passages.

**Figure 3.**
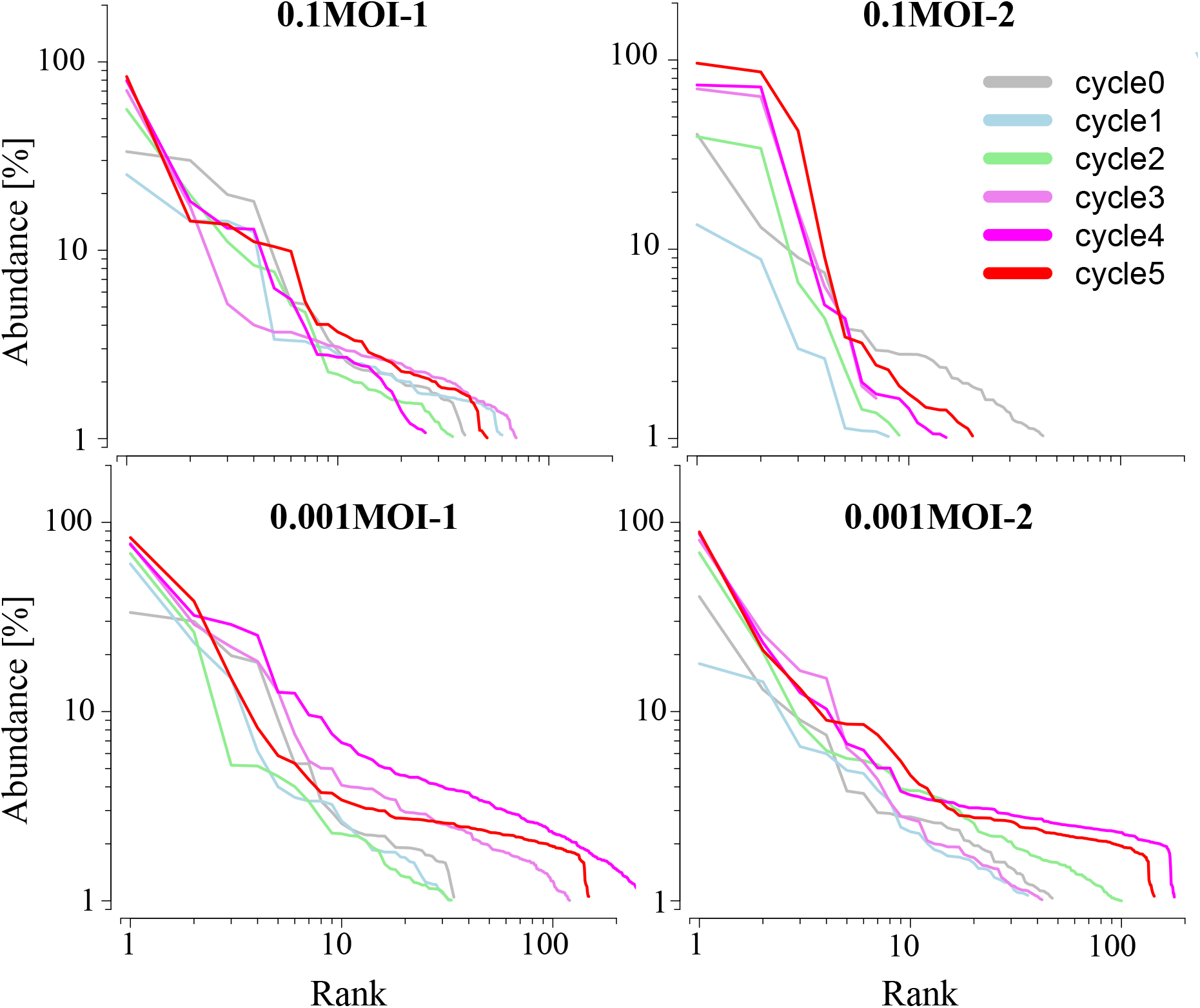
Transition of rank abundance distribution. Rank abundance distributions of each lineage are shown for every passage number (grey: initial, light blue: cycle 1, green: cycle 2, light pink: cycle 3, magenta: cycle 4, and red: cycle 5). The hemline of 0.001 MOI lineages spread and the slope became flatter as the passage repeated.

### The transition of SNPs frequency

SNPs among the entire genomic region detected using FLDS method in this study and its frequency (base 2 logarithm) were shown as a heatmap of a circos plot (Figure 4). Compared to the 0.1MOI lineages, relatively more SNPs were observed in 0.001MOI lineages. The frequency of several SNPs of 0.001MOI lineages increased after cycle 4. On the other hand, several SNPs suddenly (or randomly) appeared in or disappeared from a population. Two replacements on consensus sequences were observed VP4:V184A (common to all lineages) and VP7:E256G (specific to the 0.1MOI-2 lineage) gradually increased and finally fixed in the population (Table 1).

**Figure 4.**
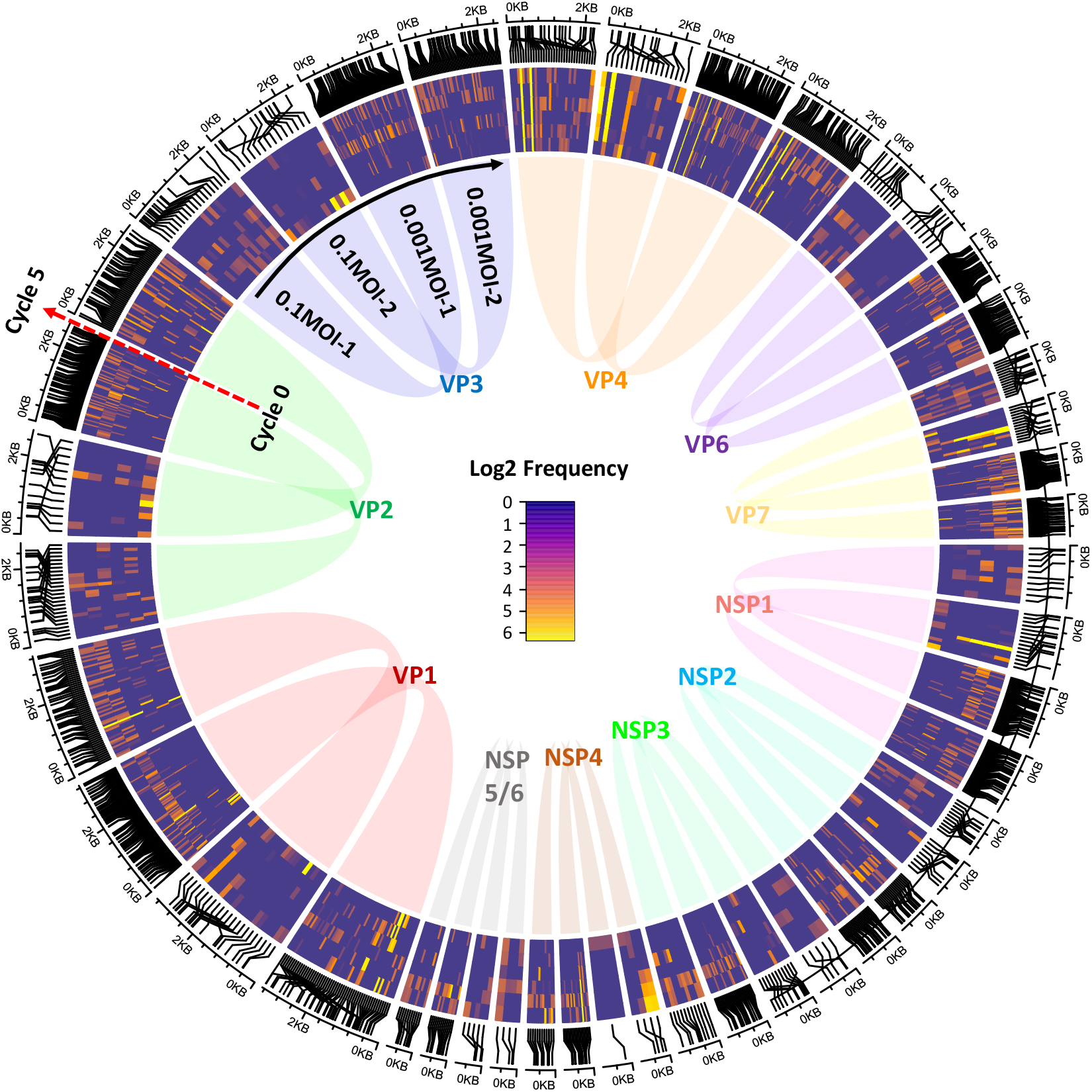
Circos plot of heatmaps of SNPs in four lineages and its frequency. Frequency of each SNP was transformed to the base 2 logarithm corresponding to the tint (high frequency: yellow, low frequency: navy). Moving from the inside to the outside of a circos plot, the log2 frequency of SNPs observed in initial population can be seen (red arrow). A heatmap block of four lineages was arranged clockwise from VP1 to NSP5/6.

**Table 1.**
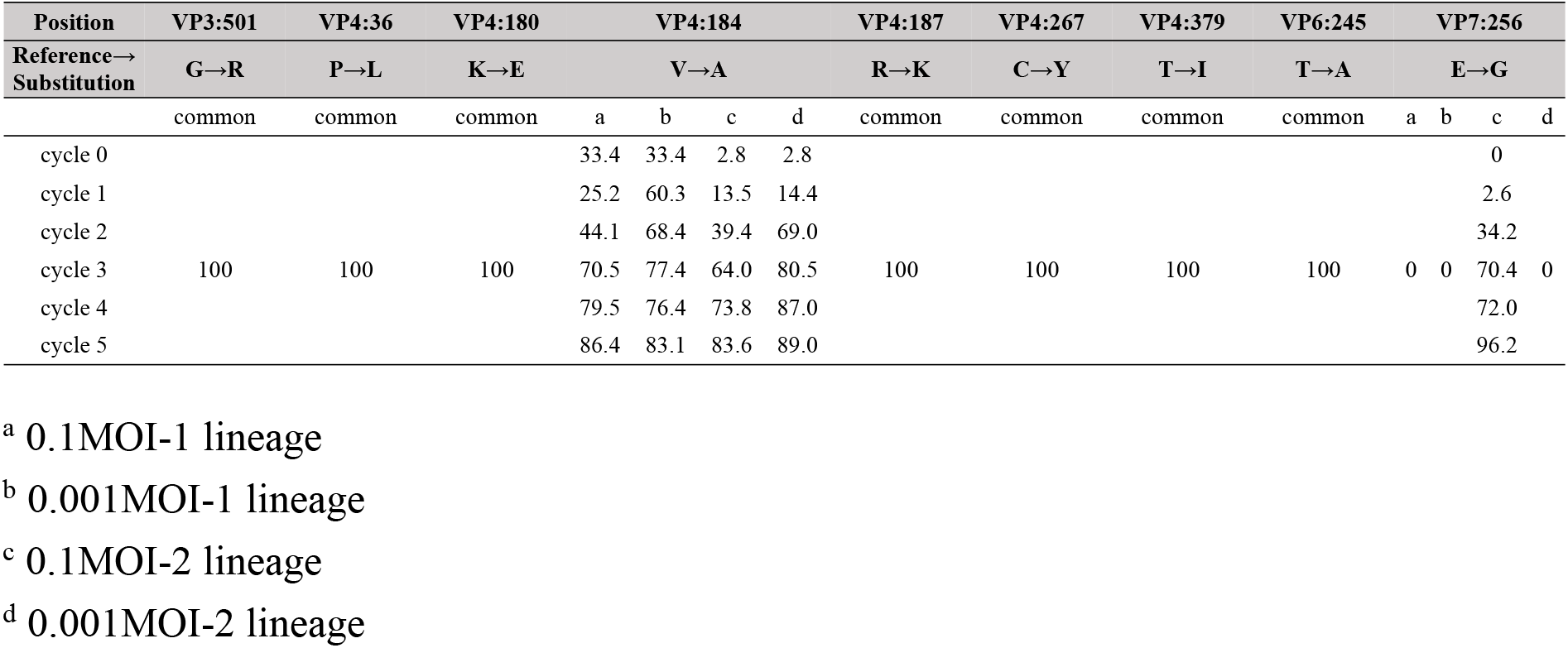
Amino acid difference between the reference and consensus sequence obtained in this study (values showed a frequency of sequences possessing a substitution [%])

### Non-synonymous and synonymous substitutions

The number of non-synonymous substitutions of the 0.001MOI lineages tends to be higher than 0.1 MOI lineages (Figure 5). The synonymous substitution rates were higher than those of non-synonymous substitutions, whereas in the case of the VP4 genome segment the rates of non-synonymous substitutions were comparable to those of synonymous substitutions. The substitutions of all genome segments occurred randomly, although spike head (about 65-224 amino acid sequence) and antigen domain (about 250-480) regions of the VP4 genome segment showed high variability.

**Figure 5.**
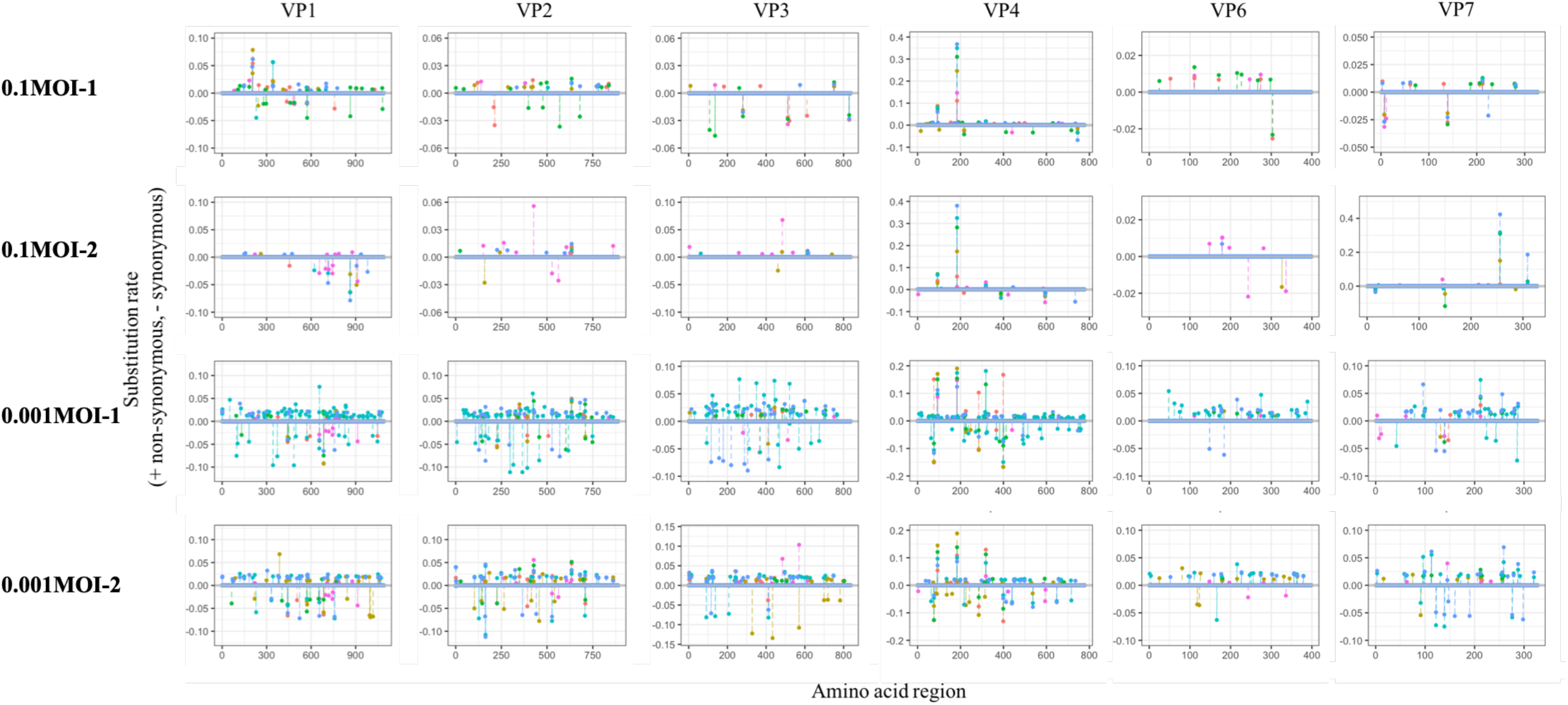
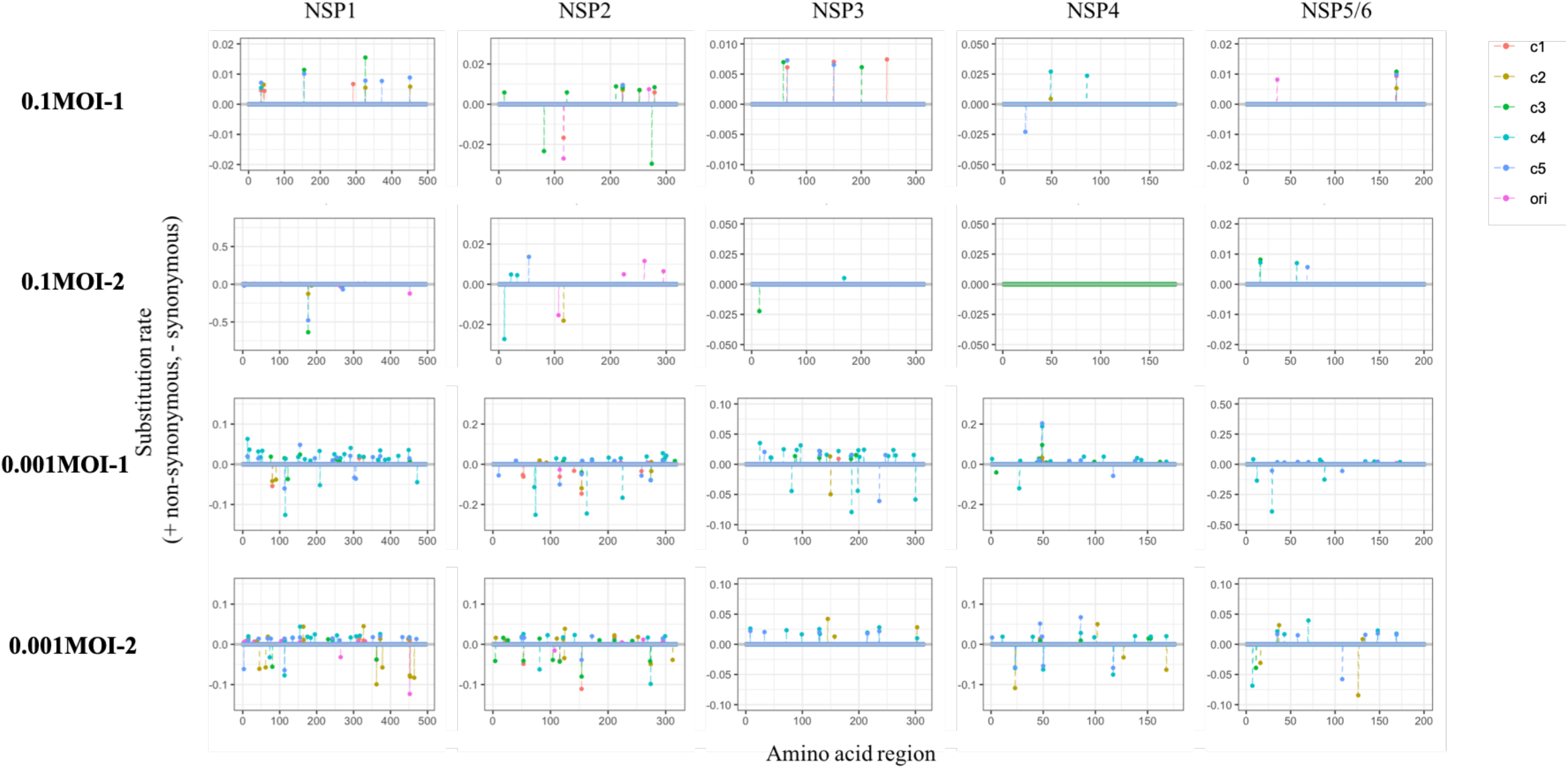
Location of the non-synonymous and synonymous substitutions. The non-synonymous substitution rate is expressed as positive, whereas the synonymous rate is shown as a negative value on each amino acid sequence in this study (pink: initial, orange: cycle 1, dark yellow: cycle 2, green: cycle 3, cyan: cycle 4, blue: cycle 5 population). **a**: Genome segments coding viral proteins. **b**: Genome segments coding non-structural proteins.

### Transition of the dN/dS ratio

The dN/dS ratio of the VP1 genome segment of the 0.001MOI lineages remained at unity, although the ratios of the 0.1MOI lineages became slightly dispersed (Figure 6). The dN/dS ratios of VP2-4, NSP1, 2, 4 and 5/6 genome segments of the 0.001MOI lineages were also around unity, but their substitution rate increased. The dN/dS ratio tended to be on the line of dN = 0 in NSP genome segments of the 0.1MOI lineages.

**Figure 6.**
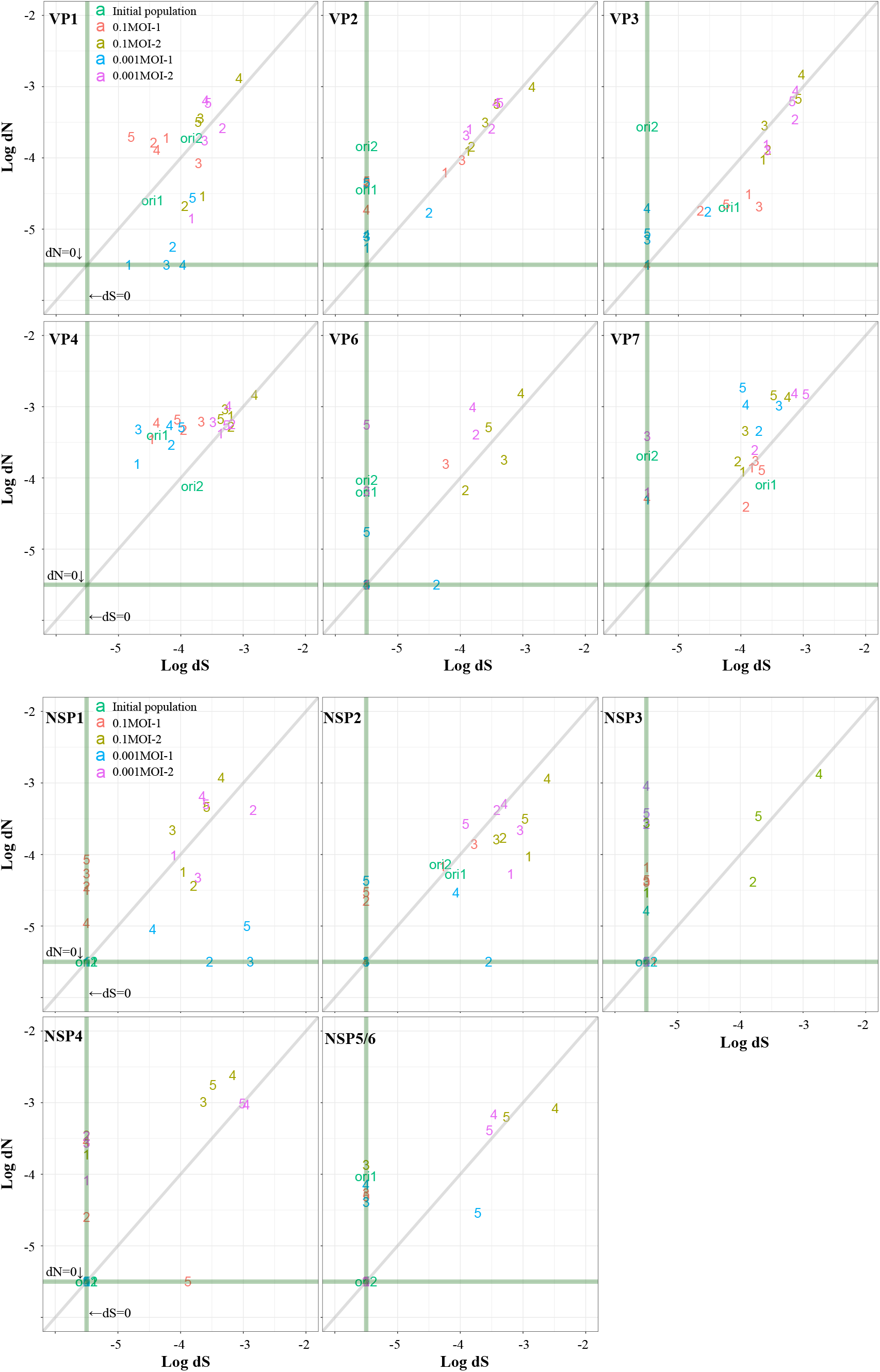
Overall proportion of the non-synonymous (dN) and synonymous (dS) substitutions in each genome segment. The proportion of logarithmic non-synonymous and synonymous substitution rate is displayed (ori1: initial population of 1st lineage, ori2: initial population of 2nd lineage, orange: 0.1 MOI-1, dark yellow: 0.001 MOI-1, blue: 0.1 MOI-2, pink: 0.001 MOI-2). The grey line indicates that the proportion of dN and dS is unity. If dN or dS is equal to zero; plots are on the green lines.

### Mutation rate of 11 genome segments per cycle

The mutation rate of each genome segments was calculated using the BEAST2 software (Table 1). Shorter genome segments such as NSP4 (750 bp) and NSP5/6 (667bp) displayed a relatively high mutation rate (about 10^−4^ substitutions/site/cycle). On the other hand, VP1 (3267 bp), VP2 (2664 bp), VP3 (2508 bp) and VP4 (2362 bp) genome segments showed an approximate 10^−5^ substitutions/site/cycle. Middle-sized genome segments (VP6, VP7, NSP1, NSP2 and NSP3) were different between the MOI settings. The NSP4 mutation rate of the 0.1MOI-2 lineage could not be calculated because no SNPs were detected.

### Simulation of the effect of minor mutants on a specific growth rate

In *Simulations 1* and *4*, the specific growth rate as a population (0.53 and 0.56 [h^−1^], respectively) was close to the value of the dominant strain (Figure 7). When the frequency of minor mutants increased to 10% (*Simulation 2*), the specific growth rate as a population increased (0.79 [h^−1^]). In *Simulation 3*, the specific growth rate of a population reached 1.03 [h^−1^], which was close to the observed value of 0.001 MOI-1_5 population (1.08 [h^−1^]).

**Figure 7.**
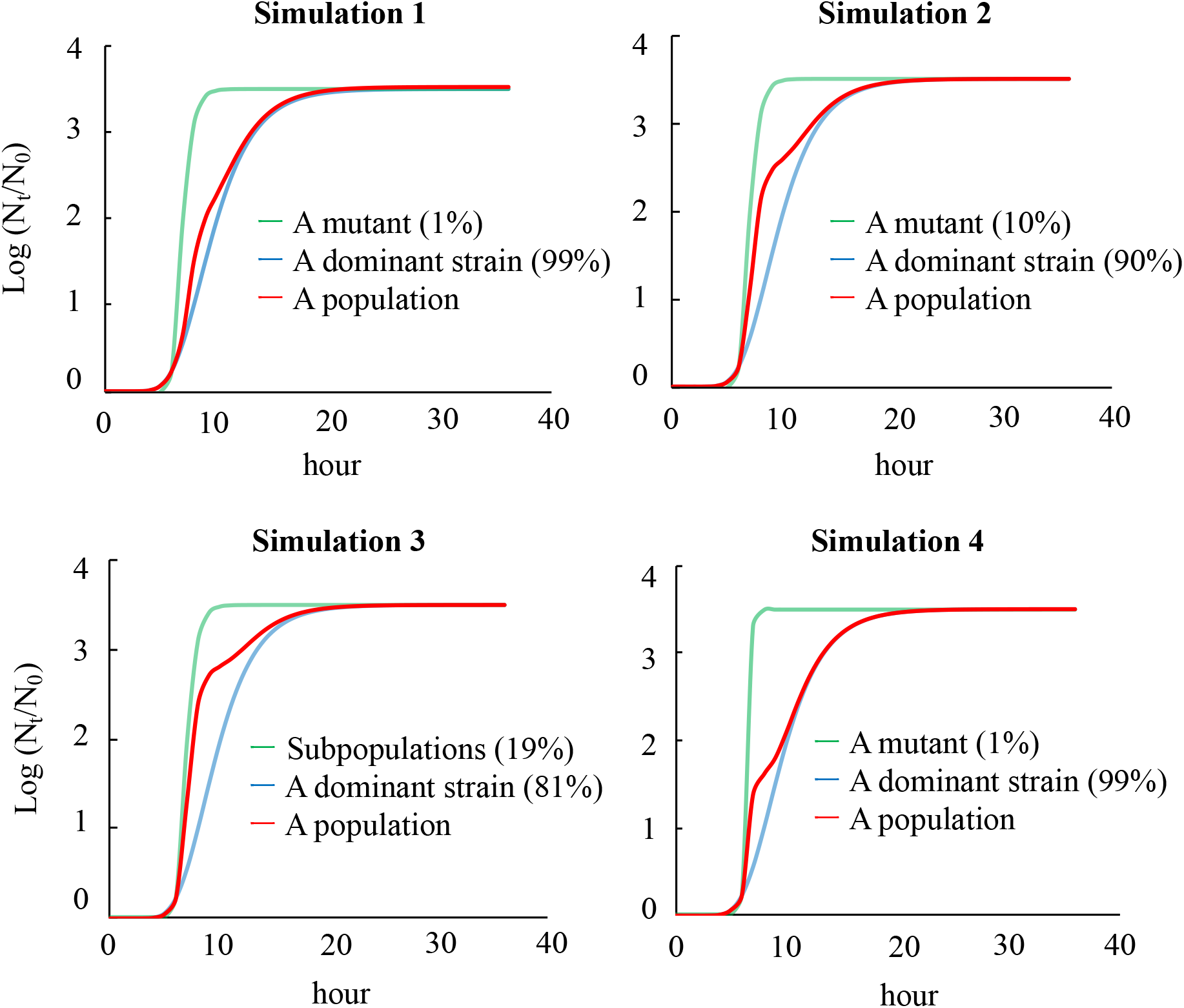
Simulation of the effect of minor mutants on the specific growth rate of a population. *Simulation 1*: the frequency of a dominant strain was 99 %, whereas that of minor mutants was 1 %. The specific growth rate of a dominant strain was set as 0.5 (h^−1^) (common among all four scenarios), but that of minor mutants was set as 2.0 (h^−1^). *Simulation 2*: the frequency of minor mutants was set as 10 %. *Simulation 3:* we assumed that 19 independent subpopulations possessing each nucleotide replacement were in a population. The frequency of the 19 subpopulations was 1.0 % each (in total 19%). *Simulation 4:* the frequency of minor mutant was 1 %, but the specific growth rate was 5.0 (h^−1^).

## DISCUSSION

In this study, RRV populations were serially passaged under stronger/weaker bottleneck conditions. The specific growth rate and the nucleotide diversity of 0.001MOI lineages (passaged under a lower MOI) increased during the latter part of serial passages (Figures 1c and 2). The mutant swarms of 0.001MOI lineages were expanded as shown in Figure 3, and a number of SNPs randomly appeared in or disappeared from RRV populations (Figure 4). Almost all mutations of 0.001MOI lineages were selectively neutral since synonymous and non-synonymous substitutions were non-site-specifically located on genome segments (Figure 4 and 5), and the dN/dS ratio was estimated to be almost unity (Figure 6). The mutation rates of each genome segment were estimated based on the SNPs information (Table 2), and the values of 0.001MOI lineages were relatively lower. Simple simulations based on the modified Gompertz model showed that minor mutants could affect the phenotype of the population in the mutant frequency-dependent manner (Figure 7).

**Table 2.** Mutation rate for rotavirus genome segments at nucleotide level.

|  | Mutation rate<sup>a</sup> (95%HPD)<sup>b</sup> |  |  |  |  |  |  |  |  |  |  |
| --- | --- | --- | --- | --- | --- | --- | --- | --- | --- | --- | --- |
|  | VP1<br>[3267 bp]<sup>c</sup> | VP2<br>[2664 bp] | VP3<br>[2508 bp] | VP4<br>[2362 bp] | VP6<br>[1356 bp] | VP7<br>[1062 bp] | NSP1<br>[1599 bp] | NSP2<br>[1059 bp] | NSP3<br>[1078 bp] | NSP4<br>[750 bp] | NSP5<br>[667 bp] |
| 0.1MOI-1 | 2.45<br>(0.05-5.69) | 5.47<br>(0.09-11.9) | 7.08<br>(0.01-17.0) | 4.55<br>(0.06-10.9) | 19.2<br>(3.76-44.8) | 16.5<br>(0.04-41.2) | 14.7<br>(0.01-36.7) | 25.3<br>(0.07-55.7) | 58.4<br>(2.22-38.9) | 44.4<br>(1.91-122) | 23.0<br>(0.20-61.2) |
| 0.1MOI-2 | 3.60<br>(0.01-8.38) | 5.32<br>(0.01-13.3) | 6.66<br>(0.02-16.5) | 5.10<br>(0.01-19.4) | 57.2<br>(0.91-67.1) | 12.8<br>(0.01-32.1) | 11.0<br>(0.02-32.1) | 19.7<br>(0.08-46.9) | 29.6<br>(0.34-82.5) | - | 27.2<br>(1.56-68.0) |
| 0.001MOI-1 | 1.56<br>(0.02-13.0) | 1.32<br>(0.03-5.30) | 0.70<br>(0.01-7.56) | 2.46<br>(0.04-39.5) | 3.48<br>(0.02-20.3) | 6.80<br>(0.05-15.8) | 6.28<br>(0.02-30.2) | 4.49<br>(1.14-14.0) | 12.0<br>(0.89-28.9) | 22.0<br>(3.38-46.3) | 4.93<br>(0.01-15.5) |
| 0.001MOI-2 | 1.68<br>(0.07-4.20) | 1.94<br>(0.03-5.16) | 2.20<br>(0.03-5.72) | 1.69<br>(0.04-3.95) | 5.95<br>(0.03-14.5) | 8.56<br>(0.01-18.8) | 7.29<br>(0.02-28.9) | 6.67<br>(0.01-19.0) | 18.9<br>(0.04-42.7) | 25.6<br>(0.01-56.0) | 21.2<br>(0.03-53.1) |

<sup>a</sup> 10<sup>-5</sup> substitutions/nucleotide site/cycle
<sup>b</sup> 95% highest posterior density
<sup>c</sup> genome length shown as base pair

A bottleneck effect randomly eliminates genome sequences from a population (31) and promotes the evolution of the virus due to a drastic change of population structure. In this study, the nucleotide diversity of rotaviruses passaged under the stronger bottleneck effect increased more than that under the weaker bottleneck effect (Figure 2). Lequime *et al.* reported that the genetic diversity of dengue virus during intra-host transmission was immediately replenished following a bottleneck event, although its expansion of a mutant swarm was also restricted by a negative selection (6). Morelli *et al.* reported that many minor mutants were transmitted during inter-host transmission where a bottleneck effect worked (32). Taken together with our study, although a bottleneck effect temporarily decreases the population size, it can excavate minor sequences from a mutant swarm. Then, the size of a mutant swarm can be expanded to the level determined by the degree of heterogeneity of the population. In our study, we determined that minor sequences in the 0.001MOI lineages found new spaces previously dominated by master sequences, and the size of mutant swarms continued to expand.

The concept of sequence space (33) provides a more detailed clue to address the question of why the increase of genetic diversity occurred in the lineages of viruses placed under the stronger bottleneck conditions. When a virus population undergoes a bottleneck, several minor sequences in a hidden layer of the mutant swarm, which originally cannot out-compete the master sequence, accidentally appear by moving into a new sequence space. In the concept of fitness landscapes (higher fitness strains are on the top of the hill, and lower fitness strains gather around the bottom of the hill) (5), the top of the hill is truncated by a bottleneck effect and then the substratum is expanded. In other words, the hill becomes less steep because the minor sequences originally existing in the hidden layer of a mutant swarm acquire some of the spaces of the upper layer of the fitness hill. Lauring and Andino suggested that mutants tend to gather in the upper part of the fitness hill if they show a low mutation rate (34), which is consistent with our results for mutation rates (Table 2). A bottleneck effect gives a chance to minor sequences of mutants to explore a sequence space.

An expansion of mutant swarms appears to lead to an increase in the specific growth rate without minor mutants being dominant in the population. Serial passages are known to change viral phenotypes. For example, the rabies virus adapted to a new environment without replacement of the master sequence during serial passages (4). An infectivity of simian/human immunodeficiency virus increased after serial passages on macaques as the size of the mutant swarm expanded (35). A replicative fitness of hepatitis C virus increased by serial passages under a bottleneck according to an expansion of the mutant swarm (9). It was also reported that the size of the mutant swarm positively correlated with the growth ability of the West Nile virus (3). These previous reports indicate that a larger mutant swarm can provide a virus population with the chance to acquire some advantageous phenotypes, which is also the case in the present study, in which the specific growth rate (relating to the growth ability) increased under a bottleneck effect (Figure 1c) as the size of the mutant swarm of the 0.001MOI lineages expanded (Figure 3). We deduce that some minor sequences find new sequence spaces created by a bottleneck, resulting in an increase in the specific growth rate as a population. Fitness is an overall indicator of the ability to produce infectious progeny, such as a binding efficiency to receptors, specific growth rate and burst size (33). The fitness of mutants displaying a higher specific growth rate was lower in this study since nucleotide replacements specific to 0.001 MOI lineages on master sequences were not found (Table 1). If the fitness of a mutant is outstanding, the frequency of the mutant in a population increases through passages and finally becomes dominant. Since many mutations were regarded as neutral in this study (Figure 5) despite an increment in specific growth rate, we speculate that several minor mutants showed inferior aspects of the fitness to the master sequences in compensation for higher growth ability.

Four simple simulations using the modified Gompertz model gave us some insight into the mechanisms of minor mutants (or small subpopulations in a mutant swarm) associated with a viral phenotype as a population (Figure 7). We found an increase in specific growth rate depending on the frequency of minor mutants or several subpopulations. From these simulations, we gained two insights about the process. First, when the frequency of one subpopulation with a higher specific growth rate is more than 10 %, the phenotype (growth rate) of the population can be changed. Second, in spite of lower frequency, several subpopulations independently affect the viral phenotype (growth rate) of a population. These findings need to be verified in a future study using isolated minor mutants with a higher specific growth rate and measuring the specific growth rate of a population.

Rotavirus outbreaks are still being reported even after implementation of vaccine programs. Some cases of rotavirus in California were reported in spite of a vaccination program and hygienic interventions such as hand washing and disinfection (13). Zeller *et al.* found that vaccination did not affect rotavirus population size, although some unique clusters were found after vaccination (36). Therefore, rotavirus persistence in human society should be explained by not only the introduction of a vaccine but also other factors. A bottleneck effect can often be seen in a life cycle of rotaviruses in a real environment (33). Human-to-human transmission corresponds to this situation of a bottleneck effect, since a few virions attached to hands can transmit to new susceptible humans from feces. Interventions such as those in California may act as a bottleneck effect for rotavirus. Although the information related to the size of mutant swarms of rotavirus populations is rarely obtained from outbreak cases, to understand the epidemic pattern of rotaviruses, we need to explore the transition of a genetic population structure of an outbreak-associated strain with molecular epidemiological studies using next generation sequencing.

## ACKNOWLEDGMENT

This study was supported by JSPS KAKENHI Grant Number 15H02272 and 18H05368, and MEXT KAKENHI Grant Number 16H06429, 16K21723, and 16H06437. We thank Prof. Keita Matsuno, Faculty of Veterinary Medicine, Hokkaido University, for his experimental support.

